# MolClustPy: A Python Package to Characterize Multivalent Biomolecular Clusters

**DOI:** 10.1101/2023.03.14.532640

**Authors:** Aniruddha Chattaraj, Indivar Nalagandla, Leslie M. Loew, Michael L Blinov

## Abstract

**Summary:** Low-affinity interactions among multivalent biomolecules may lead to the formation of molecular complexes that undergo phase transitions to become extra-large clusters. Characterizing the physical properties of these clusters is important in recent biophysical research. Due to weak interactions such clusters are highly stochastic, demonstrating a wide range of sizes and compositions. We have developed a Python package to perform multiple stochastic simulation runs using NFsim (Network-Free stochastic simulator), characterize and visualize the distribution of cluster sizes, molecular composition, and bonds across molecular clusters and individual molecules of different types.

**Availability and implementation:** The software is implemented in Python. A detailed Jupyter notebook is provided to enable convenient running. Code, user guide and examples are freely available at https://molclustpy.github.io/

**Supplementary information:** Available at https://molclustpy.github.io/

## 1 Introduction

Clustering of weakly interacting multivalent proteins and nucleic acids leads to biomolecular condensate formation via phase transition [1, 2]. These condensates are membrane-less sub-cellular compartments that play an important role in spatiotemporal regulation of cellular biochemistry [3]. Dysregulation of condensate biology is implicated in a series of pathological conditions [4-6].

Since clustering of multivalent biomolecules underlies the condensate formation, it is important to model and characterize cluster formation. For example, size and composition of the condensates have important consequences in cell signaling [7, 8]. A better understanding of the biophysical properties of such condensates may facilitate testing hypotheses, interpreting experimental observations, and developing strategies to modulate such systems in a controlled manner, potentially leading to identification of drug targets.

Molecular clustering shows a switch-like behavior (phase transition) in a concentration dependent manner [9]. Below a threshold concentration, molecules remain in the monomeric and small oligomeric states (Fig 1A, dispersed state) that will manifest as a single homogeneous phase. Upon crossing the threshold, the system tends to form large clusters (Fig 1A, clustered state). The co-existence of large clusters along with small clusters results in a splitting of the system into two distinct phases (dense and dilute). The cluster size distribution would shift from an exponentially decaying unimodal distribution (Fig 1B, left) to a bimodal (bifurcated) distribution (Fig 1B, right).

**Fig. 1.**
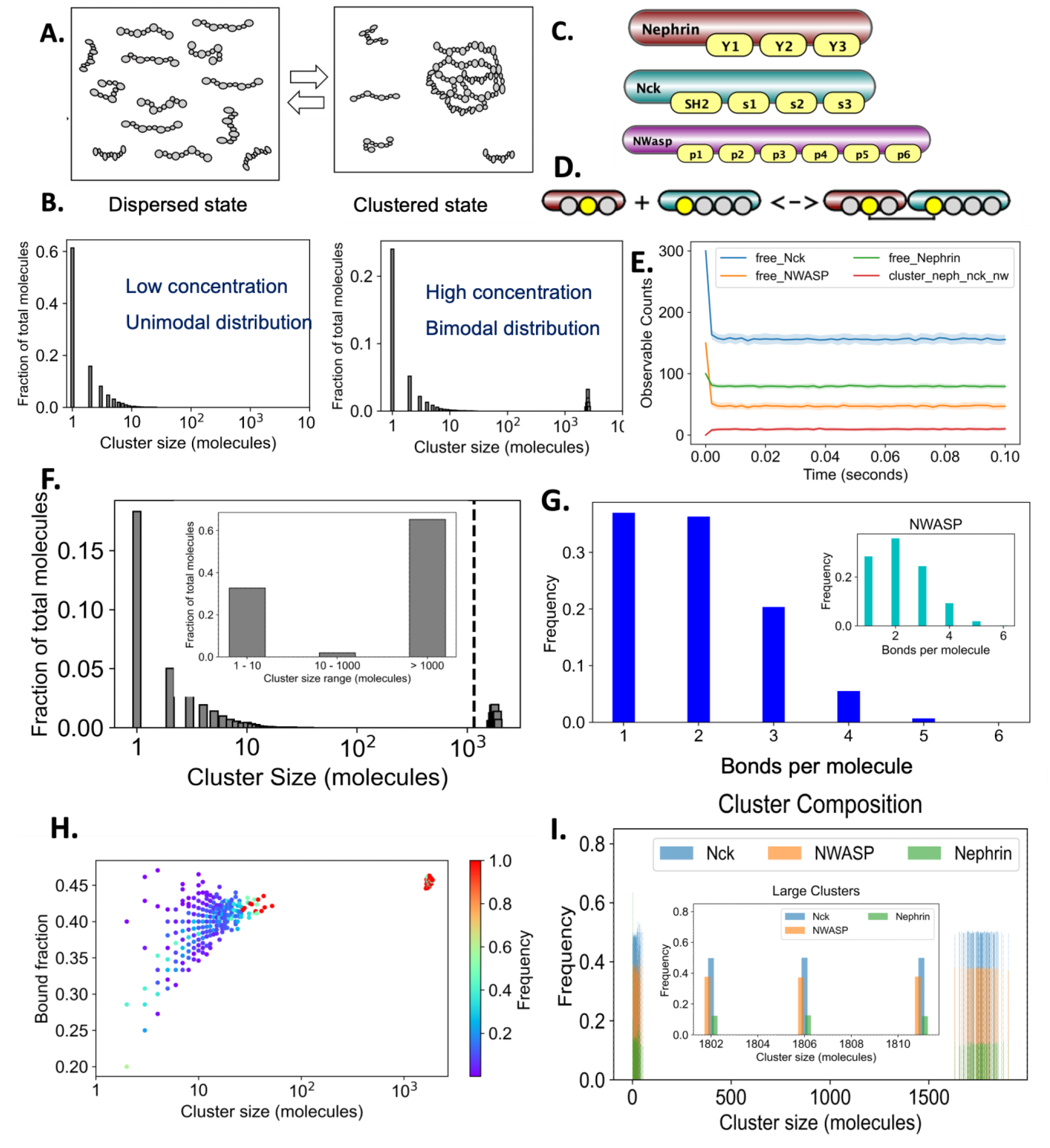
Characterization of molecular clusters. (A) Two states of molecular clusters: dispersed (monomeric and small oligomeric molecular complexes) and clustered (large clusters along with small clusters). (B) Quantification of cluster size distributions, with dispersed state being unimodal and clustered state being bimodal. (C) Rule-based depiction of three multi-site molecules Nephrin (three phosphotyrosines), Nck (one SH2 and three SH3 domains) and NWASP (six PRM domains). (D) A rule of Nephrin-Nck binding. Sites in grey do not affect the outcome of interaction, so they can be bound or unbound. (E) Simulation output for several observables, illustrating the envelope across multiple trials. (F) Average Cluster Occupancy (ACO), fraction of molecules in clusters of a given size. In insert is binning - the fractions of molecules in clusters of certain size range. (G) The average number of bonds per molecule – across all molecules and for NWASP in the insert. (H) The distribution of binding saturations across molecular clusters of different size. (I) Composition of clusters, i.e., relative abundance of each molecular types across cluster sizes. Inset shows compositions of a few large clusters.

Weak molecular interactions result in a distribution of size and compositions of multi-molecular clusters [10]. Modeling this process requires multiple stochastic simulations and subsequent statistical analysis. We have previously used *ad hoc* code to analyze multivalent clustering for different biological systems [9, 11-13]. In this work, we present a Python package – MolClustPy that generalizes these methods to statistically analyze and visualize molecular clusters from multiple stochastic trials.

The user needs to describe the molecules and their interactions in the rule-based BioNetGen Language (BNGL) format [14, 15]. Then MolClustPy will utilize the existing Python wrapper of BioNetGen (pyBioNetGen) to simulate the cluster formation for a user-defined number of times. The collected statistics averaged over multiple runs is used to provide average time courses of certain user-defined quantities (“observables”), as well as characterization of the final state of the system.

## 2 Characterization of cluster composition

### 2.1 Biological system specification and simulation

Because of stochastic nature of biomolecular condensates and very flexible molecular compositions and connectivity, stochastic agent-based modeling is the most appropriate for providing a comprehensive understanding at the system level. The most convenient way to define agents that are multi-site molecules and their interactions is via a modeling technique called rule-based modeling [15].

To demonstrate the utility of our package, we will consider an experimentally well-characterized multivalent system - Nephrin, Nck and N-WASP [16, 17]. Molecules are defined as objects with multiple binding sites. (Fig. 1C), and the interactions between molecules are defined as interactions leading to formation or breaking a bond between specific binding sites. Such interactions are best described by rules that define the input and output of interaction depending on the initial state of all binding sites. The rule in Fig 1D defines binding of Nephrin Y2 site to Nck SH2 site (sites shown in yellow). This rule corresponds to a potentially infinite number of individual interactions among molecular clusters, because other sites of interaction molecules (shown in grey) may be bound to other molecules not shown in the rule definition. However, all interactions corresponding to the single rule are parameterized by the same on- and off-rate constants. Note that rules are not limited to binary site-site interactions but can be extended to encode arbitrary levels of complexities for interaction among multi-valent molecules; for example, the binding strength of one molecular site may depend on the binding status of other sites of the same molecule to capture allosteric or cooperativity effects. Once two molecules are part of a complex, an additional bond (intra-complex) formation may be prevented (zero affinity) or amplified (higher affinity) to avoid or promote ring formation.

To stochastically simulate rule-based systems with a potentially infinite number of species and interactions we use an agent-based Network-Free stochastic simulator NFsim [18]. NFSim simulation outputs consist of two major parts – observables and final molecular configuration. Observables are predefined global properties of the biological system whose concentrations are reported as a time course during the simulation, for example, concentrations of free molecules of a certain type. The final molecular configuration is the set of molecular complexes that exist at the end of single simulation run.

### 2.2 MolClustPy outputs

MolClustPy analyzes both observables and molecular clusters across multiple simulation runs. Figure 1E illustrates plots of the time courses for observables. Importantly, we demonstrate both the mean trajectory and the standard deviation shown as a fluctuation envelope. The larger is system’s stochasticity, the wider is the envelope. To demonstrate the width of the distribution, we plot observables for a lower concentrations of molecules.

Figure 1F illustrates the cluster size distribution: each bar corresponds to the fraction of total molecules in each cluster size. In other words, this is a probability distribution of finding molecules in a certain cluster size. For example, the probability of finding a molecule in a monomeric form is 17%. The mean of the distribution is called average cluster occupancy (ACO) as shown by the dashed line. For a large cluster size range, a binned histogram might be helpful, as shown in the inset. Here we see that 33% of all molecules are in small clusters of sizes up to 10, while around 62% of molecules are in large clusters (> 1000 molecules).

Figure 1G captures the molecular crosslinking – the average number of bonds per molecule. For our mixed-valent system, Nephrin can have 1-3 bonds, Nck can have 1-4 bonds, while NWASP may have 1-6 bonds. In that case, inspecting bond distribution of individual molecular types (NWASP, inset) gives more intuitions.

Figure 1H demonstrates the degree of bond saturation across molecular clusters of different sizes. Bound fraction (BF) is the ratio of bound sites over total sites present in that cluster. The color bar shows the relative frequencies of a given configuration. From the BF pattern we see that there is a greater variability in smaller clusters, while large clusters converge to a fixed BF likely due to entropic reasons.

Figure 1I demonstrates the molecular composition of the clusters, giving the relative fraction of each molecular type within a given cluster size. Note that within each bar the sum of all fractions is 1. The large clusters seem to have identical compositions (inset), suggesting a stoichiometry driven clustering process which is consistent with recent experimental finding [8].

## 3 Implementation

The package is implemented in Python. It requires the pyBioNetGen package to run. MultiRun_BNG.py processes the BNGL file that defines molecules, initial species, rules of interactions and optional observables. The user must input three parameters related to simulation: duration of each simulation, number of output time points, and number of stochastic trials. MultiRun_BNG.py calls the pyBioNetGen package which executes NFsim simulations the required number of times. NFsim_data_analyzer.py collects the observables and molecular clusters to perform various statistical analyses. DataViz_NFsim.py visualizes the statistical outputs.

To quantify the cluster property distribution, we collect the molecular species (clusters) from multiple trials and perform statistical analysis on the combined dataset. A “cluster” is a molecular network where each node is a multivalent molecule, and the bonds comprise the edges. We analyze the network topology and their relative abundances. For example, the degree of a node gives the number of bonds coming out of that particular molecule. Bound fraction is then computed as the ratio of bound sites over total sites present in a molecule.

## References

1. Banani, S.F., et al., Biomolecular condensates: organizers of cellular biochemistry. Nat Rev Mol Cell Biol, 2017. 18(5): p. 285–298.

2. Choi, J.-M., A.S. Holehouse, and R.V. Pappu, Physical Principles Underlying the Complex Biology of Intracellular Phase Transitions. Annual Review of Biophysics, 2020. 49(1): p. 107–133.

3. Lyon, A.S., W.B. Peeples, and M.K. Rosen, A framework for understanding the functions of biomolecular condensates across scales. Nat Rev Mol Cell Biol, 2021. 22(3): p. 215–235.

4. Alberti, S. and D. Dormann, Liquid-Liquid Phase Separation in Disease. Annu Rev Genet, 2019. 53: p. 171–194.

5. Mathieu, C., R.V. Pappu, and J.P. Taylor, Beyond aggregation: Pathological phase transitions in neurodegenerative disease. Science, 2020. 370(6512): p. 56.

6. Wang, B., et al., Liquid-liquid phase separation in human health and diseases. Signal Transduct Target Ther, 2021. 6(1): p. 290.

7. Su, X., et al., Phase separation of signaling molecules promotes T cell receptor signal transduction. Science, 2016. 352(6285): p. 595–9.

8. Case, L.B., et al., Stoichiometry controls activity of phase-separated clusters of actin signaling proteins. Science, 2019. 363(6431): p. 1093–1097.

9. Chattaraj, A., M.L. Blinov, and L.M. Loew, The solubility product extends the buffering concept to heterotypic biomolecular condensates. eLife, 2021. 10: p. e67176.

10. Mayer, B.J., M.L. Blinov, and L.M. Loew, Molecular machines or pleiomorphic ensembles: signaling complexes revisited. Journal of Biology, 2009. 8(9): p. 81.

11. Falkenberg, C.V., M.L. Blinov, and L.M. Loew, Pleomorphic ensembles: formation of large clusters composed of weakly interacting multivalent molecules. Biophysical journal, 2013. 105(11): p. 2451–2460.

12. Falkenberg, C.V., J.H. Carson, and M.L. Blinov, Multivalent Molecules as Modulators of RNA Granule Size and Composition. Biophys J, 2017. 113(2): p. 235–245.

13. Chattaraj, A., M. Youngstrom, and L.M. Loew, The Interplay of Structural and Cellular Biophysics Controls Clustering of Multivalent Molecules. Biophys J, 2019. 116(3): p. 560–572.

14. Blinov, M.L., et al., BioNetGen: software for rule-based modeling of signal transduction based on the interactions of molecular domains. Bioinformatics, 2004. 20(17): p. 3289–91.

15. Faeder, J.R., Blinov, M. L., & Hlavacek, W. S., Rule-based modeling of biochemical systems with BioNetGen., in Systems Biology I.V. Maly, Editor. 2009, Humana Press. p. 113–167.

16. Li, P., et al., Phase transitions in the assembly of multivalent signalling proteins. Nature, 2012. 483(7389): p. 336–40.

17. Banjade, S. and M.K. Rosen, Phase transitions of multivalent proteins can promote clustering of membrane receptors. Elife, 2014. 3.

18. Sneddon, M.W., J.R. Faeder, and T. Emonet, Efficient modeling, simulation and coarse-graining of biological complexity with NFsim. Nat Methods, 2011. 8(2): p. 177–83.

